# Neural contributions to reduced fluid intelligence across the adult lifespan

**DOI:** 10.1101/2022.07.27.501673

**Authors:** Daniel J. Mitchell, Alexa L. S. Mousley, Meredith A. Shafto, Cam-CAN, John Duncan

## Abstract

Fluid intelligence – the ability to solve novel, complex problems – declines steeply during healthy human aging. Using functional magnetic resonance imaging (fMRI), fluid intelligence has been repeatedly associated with activation of a frontoparietal brain network, and focal damage to these regions suggests that fluid intelligence depends on their integrity. It is therefore possible that age-related functional differences in frontoparietal activity contribute to the reduction in fluid intelligence. This paper reports on analysis of the Cambridge Centre for Ageing and Neuroscience (Cam-CAN) data, a large, population-based, healthy, adult lifespan cohort. The data support a model in which age-related differences in fluid intelligence are partially mediated by the responsiveness of frontoparietal regions to novel problem-solving. We first replicate a prior finding of such mediation using an independent sample. We then precisely localise the mediating brain regions, and show that mediation is specifically associated with voxels most activated by cognitive demand, but not with voxels suppressed by cognitive demand. We quantify the robustness of this result to potential unmodelled confounders, and estimate the causal direction of the effects. Finally, exploratory analyses suggest that neural mediation of age-related differences in fluid intelligence is moderated by the variety of regular physical activities, more reliably than by their frequency or duration. An additional moderating role of the variety of non-physical activities emerged when controlling for head motion. A better understanding of the mechanisms that link healthy aging with lower fluid intelligence may suggest strategies for mitigating such decline.

**Significance Statement:** Global populations are living longer, driving urgency to understand age-related cognitive declines. Fluid intelligence is of prime importance, because it reflects performance across many domains, and declines especially steeply during healthy aging. Despite consensus that fluid intelligence is associated with particular frontoparietal brain regions, little research has investigated suggestions that under-responsiveness of these regions mediates age-related decline. We replicate a recent demonstration of such mediation, showing specific association with brain regions most activated by cognitive demand, and robustness to moderate confounding by unmodelled variables. By showing that this mediation model is moderated by the variety of regular physical activities, more reliably than by their frequency or duration, we identify a potential modifiable lifestyle factor that may help promote successful aging.

## Introduction

Aging affects many cognitive abilities, but the drop in fluid intelligence – the ability to solve novel, complex problems (Cattell, 1963; Carpenter et al., 1990; Duncan et al., 2017) – is particularly steep (Horn and Cattell, 1967; Kievit et al., 2014; Samu et al., 2017). Moreover, fluid intelligence may be especially important for determining aging trajectories through contributions to ability across cognitive domains (Parkin and Java, 1999; Salthouse et al., 2003), performance in daily life (Diehl et al., 1995), and even health outcomes (Gottfredson and Deary, 2004). Understanding fluid intelligence decline is increasingly urgent as lifespans increase (Vaupel, 2010).

Extensive research associates fluid intelligence with a frontoparietal brain network (Jung and Haier, 2007; Duncan, 2010; Santarnecchi et al., 2017a). These regions, including the precentral sulcus, middle frontal gyrus, intraparietal sulcus, anterior insula, and anterior cingulate cortex, have been termed the multiple-demand network (MDN) due to their activation during many cognitively demanding tasks (Duncan and Owen, 2000; Naghavi and Nyberg, 2005; Duncan, 2010) including tests of fluid intelligence (Prabhakaran et al., 1997; Duncan et al., 2000). Individual differences in fluid intelligence correlate with MDN activity (Gray et al., 2003; Lee et al., 2006; Tschentscher et al., 2017; Assem et al., 2020b) and with its connectivity to other brain networks (Cole et al., 2012). The impact of focal lesions (Glascher et al., 2010; Woolgar et al., 2010; Barbey et al., 2014; Woolgar et al., 2018; Smith et al., 2022) and transient transcranial stimulation (Momi et al., 2020) suggests these regions’ causal role in supporting fluid intelligence. We therefore test the possibility that functional differences in MDN activation mediate fluid intelligence decline during healthy aging (Phillips and Della Sala, 1998). Confirming this would help to understand the mechanism of age-related decline, and suggest potential targets for mitigation with interventions that might impact on the putative causal pathway from age to fluid intelligence via neural responsiveness.

Many studies have considered relations between brain activation and cognitive performance in older adults (Dennis and Cabeza, 2008; Eyler et al., 2011; Grady, 2012), although few have probed the specific three-way relationship between differences in age, brain activation and fluid intelligence. In the context of broader questions on differential age effects across task domains, a recent study (Samu et al., 2017) reported results consistent with frontoparietal activity mediating age differences in performance during a fluid intelligence task. Activity of the Default Mode Network (DMN), typically deactivated during attentionally demanding tasks (Buckner et al., 2008), was not found to mediate performance decline in the task, despite changing with age in tasks that showed behavioural decline.

Here, we have five main aims. First, we replicate the finding of frontoparietal mediation of age differences in fluid intelligence (Samu et al., 2017), using an independent, non-overlapping sample of participants from the population-based, healthy, adult lifespan cohort (Cam-CAN; Shafto et al., 2014) used by Samu and colleagues. As the dependent variable, we replace concurrent task performance with a previously-acquired, standardised fluid intelligence measure (Cattell and Cattell, 1973), avoiding external factors (e.g. arousal) that could co-modulate simultaneous measures of brain activity and behaviour. Second, after combining the non-overlapping sample with that from Samu et al. (2017), we test whether mediation is specifically associated with voxels most responsive to cognitive demands, or also with voxels suppressed by cognitive demand. Third, we assess the robustness of the mediation result to possible unmodelled covariates. Fourth, since the mediation analysis cannot itself determine causality, we estimate causal directions under additional assumptions of an acyclic model with non-Gaussian errors. Finally, in exploratory analyses, we consider whether these relationships depend on potentially modifiable lifestyle factors. Specifically, increasing research proposes that physical exercise confers resilience to cognitive aging, although the nature and mechanism of this benefit remains unclear (Colcombe and Kramer, 2003; Smith et al., 2010; Liu-Ambrose et al., 2018). We therefore examine whether the mediation model is moderated by questionnaire measures that distinguish the variety, frequency and duration of regular physical recreations.

## Materials and Methods

### Experimental Design and Statistical Analyses

The original experimental design for the Cambridge Centre for Ageing and Neuroscience (Cam-CAN) project (www.cam-can.com) is described in Shafto et al. (2014). Full details of the within- and between subject variables, statistical tests, and software used in the current project are described in the following sections. Data can be requested after registration via the Cam-CAN dataset inventory (https://camcan-archive.mrc-cbu.cam.ac.uk/dataaccess/). Analysis code is available via the Open Science Framework (https://osf.io/xgw56/). The analyses were not preregistered.

### Participants

Participants reported here are a subset of 252 participants from the population-based healthy adult cohort recruited for the Cambridge Centre for Ageing and Neuroscience project (Cam-CAN; see Shafto et al., 2014 for full details of the sample and exclusion criteria). The initial Cam-CAN data collection consisted of a background interview (Stage 1), detailed cognitive testing and core measures of brain structure and function (Stage 2), and targeted functional neuroimaging studies (Stage 3). The current study reports results from the behavioural test of fluid intelligence (Stage 2), the fMRI session using a similar fluid intelligence task (Stage 3), and self-reported recreational activities (from a questionnaire distributed in Stage 1). Of the participants recruited, 252 (133 female) completed the fMRI task, of whom all had completed the prior fluid intelligence test. Stage 3 testing occurred between 0.3 and 3.4 (mean 1.4) years following Stage 2. For analyses including age, we used the age midway between the two tests, for which ages ranged from 20.5 to 90.3 years (mean 55.1; approximately equal numbers per decile).

The first analysis, seeking to replicate the mediation observed by Samu et al. (2017), used a subset of 154 participants not included in the prior study. The remaining analyses used all 252 participants, except for the final analyses of moderated mediation, for which questionnaire data on physical recreation were missing from 13 participants and data on non-physical activities were missing from 15 participants.

Participants gave written, informed consent, and the study was conducted in accordance with ethical approval obtained from the Cambridgeshire 2 (now East of England—Cambridge Central) Research Ethics Committee.

### Fluid intelligence measure

Fluid intelligence was assessed using Scale 2, Form A of Cattell’s Culture Fair Test (Cattell and Cattell, 1973), according to the standard protocol. This consists of four, nonverbal, multiple-choice, pencil-and-paper subtests of abstract reasoning (series completion, odd-one-out, matrix completion, and topological judgement) each introduced with examples and then completed under timed conditions, but with participants not informed of the precise time limits. When the total number of correct problems was converted to its standardized fluid intelligence score (IQ) using the conversion table in the manual (which is age-adjusted only below age 14; Cattell and Cattell, 1973), its variance was found to decrease significantly with age (White-Wooldridge test, *χ*_2_(2) = 11.6, p=0.003). We therefore constructed a latent IQ variable from the first principle component across the subtests, similar to Kievit et al. (2014), which had homoscedastic residuals when predicted from age. This variable was standardised to have the same sample mean and standard deviation as the normed scores based on the manual.

### Lifestyle activities measures

Recent years have seen increasing consensus that physical exercise can be beneficial for neurocognitive health, including in older adults (Colcombe and Kramer, 2003; Smith et al., 2010; Liu-Ambrose et al., 2018). Most studies have examined a single measure of exercise, so cannot distinguish which aspects of increased exercise might be most beneficial; however some recent reports suggest that the duration of exercise may be less important than its intensity (Angevaren et al., 2007; Brown et al., 2012) or variety (Angevaren et al., 2007). In the Cam-CAN study, we had access to questionnaire measures that distinguished the variety, frequency and duration of self-reported physical recreational activities, allowing us to investigate which of these different aspects of exercise might have the greatest impact on age-related decline in fluid intelligence.

Measures of recent physical recreational activities were taken from a self-completion questionnaire, completed during Stage 1 (within 2 years of Stage 2), based on the recreation section of the EPIC-EPAQ2 questionnaire (Wareham et al., 2002), which was derived, in turn, from the Minnesota Leisure Time Activity questionnaire (Richardson et al., 1994). Questions included the approximate frequency (on a seven point scale) and duration (in hours and minutes) of each of 35 recreational and DIY activities in which people had participated over the last year (e.g. cycling, mowing the lawn, dancing, golf). We considered just “regular” activities, which we defined as those occurring at least monthly. “Variety” of regular activities was measured as the number of different activities, “frequency” per regular activity was measured as the mean number of episodes in a year, and “duration per episode” was also measured as the mean across regular activities; “total duration” was calculated by multiplying the frequency and duration of each activity and summing over regular activities. In this way, we sought to address the question: if someone were to devote a fixed amount of time to extra physical activity, might it be better to perform their current activities for longer, to perform their current activities more often, or to engage in a greater range of activities?

To assess whether the results for the variety of physical activities generalised to more intellectual activities, we derived a similar measure for the variety of recent “non-physical” activities. For these activities, duration was never reported, and for most of them frequency was not reported, so variety was the only measure. The variety of recent non-physical activities was quantified by summing the number of mental and social activities reported across two sources: (1) The “recent activities” portion of the self-completion questionnaire (based on elements of the Lifetime of Experiences Questionnaire; Valenzuela and Sachdev, 2007) provided 38 items, including mental activities in a typical week (e.g. reading, art, crossword puzzles), events or entertainment in the last two months (e.g. cinema, pub, concert), usual means of acquiring information (e.g. TV, newspapers, internet), and kinds of materials read on a regular basis (e.g. newspaper, novels, magazines); (2) from a home interview at Stage 1 we included self-reports of 22 types of social interactions (e.g. phone friends, email friends, attend social clubs). As for the physical activity measures, for all questions where frequencies were reported we only counted regular non-physical activities, i.e. those that occurred at least monthly.

### MRI acquisition

MRI was performed on a 3 Tesla Siemens TIM Trio System, using a 32 channel head coil.

A high resolution 3D T1-weighted structural image was acquired using a Magnetization Prepared Rapid Gradient Echo (MPRAGE) sequence, with the following parameters: Repetition Time (TR) = 2250 ms; Echo Time (TE) = 2.99 ms; Inversion Time (TI) = 900 ms; flip angle = 9 degrees; field of view (FOV) = 256 × 240 × 192 mm; voxel size = 1 mm isotropic; GRAPPA acceleration factor = 2.

fMRI used a T2*-weighted Gradient-Echo Echo-Planar Imaging (EPI) sequence with the following parameters: 32 axial slices (acquired in descending order); slice thickness of 3 mm, with an inter-slice gap of 25%; TR = 2 s; TE = 30 ms; flip angle = 78 degrees; FOV = 192 × 192 × 120 mm; voxel-size = 3 × 3 × 3.75 mm.

### fMRI task

During fMRI, participants performed a non-verbal reasoning task (Figure 1a) that has been previously shown to activate the frontoparietal MDN (Duncan et al., 2000; Woolgar et al., 2013) and is based on the odd-one-out subtest of Cattell’s Culture Fair test (Cattell and Cattell, 1973). The task consisted of a series of problems in which participants were presented with a horizontal display of four panels and were instructed to select the panel that differed in some way from all of the others. The horizontal extent of each display was approximately twelve degrees of visual angle. The task used a block design, with alternating blocks of easy and difficult problems. In the easy blocks, three panels were identical and the fourth was clearly different, rendering each decision trivial; in the difficult blocks, the four panels in each problem differed in many ways, requiring the identification of abstract patterns to select the odd one out. Participants completed four easy blocks and four difficult blocks, each preceded by a 3 second cue indicating whether the upcoming problems would be “Easy” or “Hard”. Each problem remained on the screen until the participant responded, whereupon the next problem was presented after a 500 ms blank interval. Problems were presented in fixed order, drawn from a pool of 320 easy and 25 difficult problems. If a participant completed all problems of a given difficulty, problems were recycled from the beginning. (Across participants, the mean percentage of repeated problems was 0.2% for easy problems, and 12.5% for difficult problems.) Each block automatically ended after 30 seconds. Participants were encouraged to puzzle over each problem for as long as necessary, only responding when confident of the correct answer. Thus the number of trials per block varied, while the time spent on each type of problem (easy and difficult) was held constant. The task was presented using E-Prime (Psychology Software Tools, Pittsburgh, PA) and stimuli were back-projected onto a screen that was viewed through a mirror mounted on the head coil. Responses were made using a button box.

**Figure 1:**
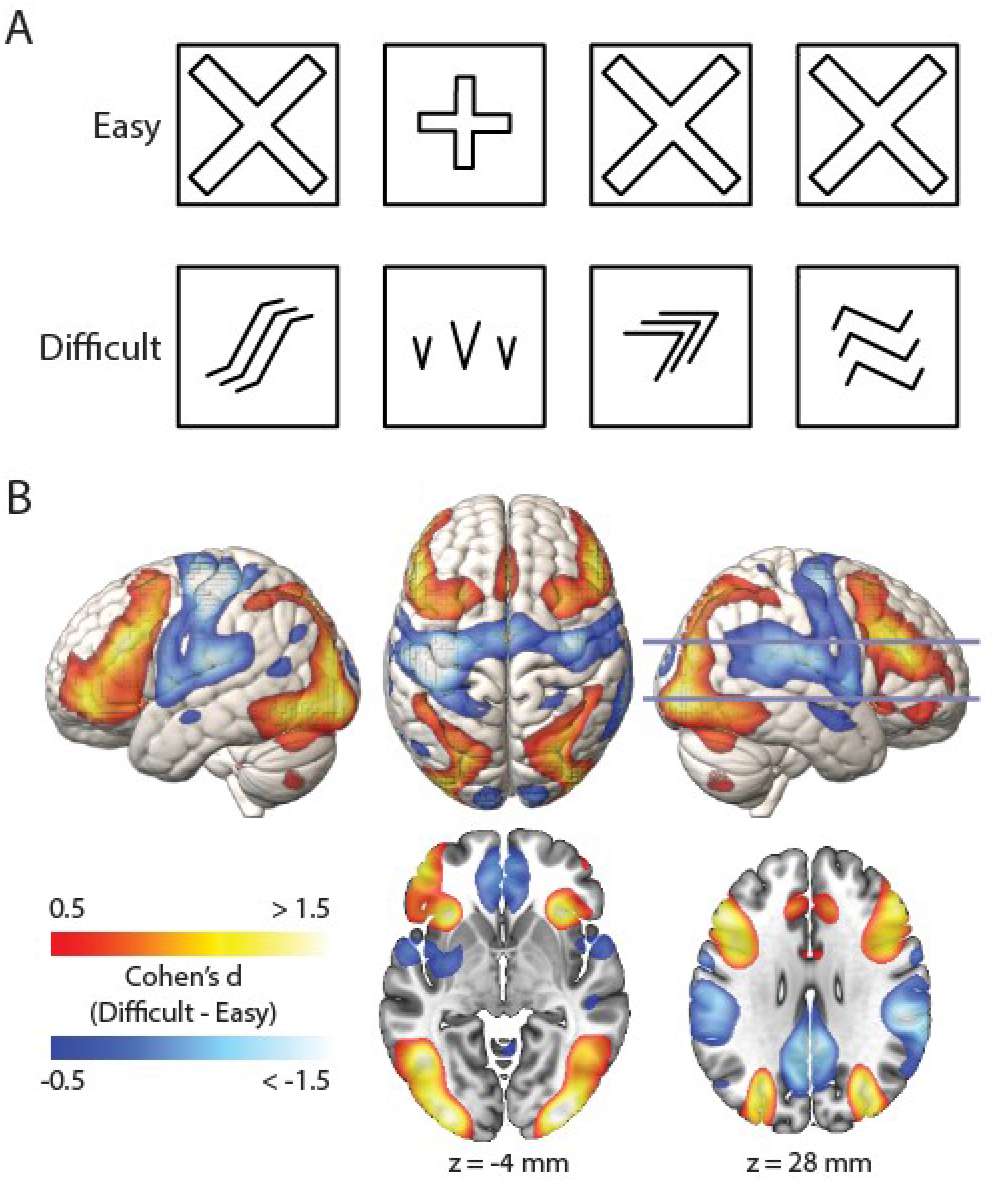
fMRI task, and response to difficult versus easy problem-solving. A) Examples of easy and difficult problems. In both examples, the 2^nd^ item is the correct answer. B) Regions significantly more active in the difficult than the easy condition (warm colours) and vice versa (cool colours), shown overlaid on surface renderings (top) and horizontal slices (bottom). N=252. Slice positions are labelled in MNI coordinates and marked on the right hemisphere rendering. Activations are shown to a depth of 15 mm. Colour scales indicate Cohen’s d, thresholded at |d| > 0.5. (All effects greater than |d| > 0.13 are significant when controlling the FDR at < 0.05.)

Before entering the scanner, participants were shown examples of the types of problems that they would encounter, and they practised selecting the odd-one-out until both they and the experimenter were happy that they understood the instructions.

### fMRI pre-processing

Data were analysed using “automatic analysis” software (Cusack et al., 2015) in Matlab (The Mathworks, Inc., Natick, MA, United States), which called relevant routines from SPM12 (Wellcome Department of Imaging Neuroscience, London, UK).

Each participant’s structural volume was segmented into probabilistic maps of six tissue classes. Grey and white matter maps of all Cam-CAN Stage 2 participants whose structural volumes passed quality control (272, including 20 participants not reported here) were non-linearly aligned using diffeomorphic registration (DARTEL; Ashburner, 2007) to create a group template volume, which was then normalised to the MNI (Montreal Neurological Institute) template via an affine transformation.

Functional volumes were rigidly realigned to correct for motion, and slice-time corrected. They were then co-registered to the structural volume, and normalised to template space using the combined transformations (native to group template, and group template to MNI template) derived from the structural volume. Functional volumes were spatially smoothed using a Gaussian kernel of 8 mm full-width-half-maximum (FWHM).

For each participant, at each voxel, a general linear model (GLM) was used to contrast the blood-oxygen-level-dependent (BOLD) response to difficult versus easy problem-solving. One regressor for each condition was constructed by convolving the duration of each block with the canonical haemodynamic response function. Additional covariates of no interest included the six movement parameters from the realignment step, and a constant regressor to model the session mean. The model and data were high-pass filtered with a cut-off of 1/128 Hz. Estimation of the model produced beta maps for each of the easy and difficult conditions. The difference map (ΔBOLD, difficult minus easy) summarised the BOLD response to difficult problem-solving for each participant. The mean group effect was assessed using a t-test per voxel.

### Regions of interest (ROIs) and voxel-wise analyses

For the analysis replicating the mediation observed by Samu et al. (2017), the fMRI response was summarised within each of twelve cortical networks reported by Ji et al. (2019), based on the multimodal parcellation from the Human Connectome Project (Glasser et al., 2016). Parcels were generated from https://identifiers.org/neurovault.image:30759, dilated to fill a grey-matter mask, and combined into one ROI per network. The core MDN regions – areas consistently and strongly activated by multiple cognitive demands – have been shown to comprise a subset of the larger frontoparietal resting state network (Assem et al., 2020a), so these regions were added as an extra “core MDN” ROI, expected to be most strongly responsive to the difficulty contrast. Similarly, a core DMN ROI was constructed using the same data and approach as in Assem et al. (2020a), but reversing the sign of the contrast (i.e. selecting parcels consistently most active in easier compared to harder conditions across a set of tasks). This identified a midline subset of the broader DMN network parcels. The response within each ROI was summarised by the mean across voxels.

Subsequent analyses used a voxel-wise approach to precisely identify those voxels where the mediation model was significant, and to assess the association between the strength of the mediation effect and the strength of the response to cognitive demand. Across most of the brain, fMRI data were available from all 252 participants; however, towards the edges of the brain data were missing from some participants depending on the position of the acquisition bounding box. We analysed all voxels within an MNI template brain mask for which fMRI data were acquired from at least 100 participants. The sample size thus ranged from 100-252 across voxels. (73% of voxels within the mask had data from all 252 participants; 90% of voxels had data from at least 90% of participants; 98% of voxels had data from more than 100 participants.)

The set of voxels exhibiting both significant mediation and significant activation by cognitive demand (and with data from all 252 participants) then served as a functional ROI across which the BOLD response was averaged to fit a summary mediation model. This summary model was used in further analyses to examine robustness to unmodelled confounders, estimated causal direction, and moderated mediation.

For both ROI-based and voxel-based analyses, multiple comparisons were accounted for by controlling the false discovery rate (FDR) at < 0.05 (Benjamini and Yekutieli, 2001). Brain renderings are displayed using MRIcroGL software (https://www.nitrc.org/projects/mricrogl).

### Mediation analyses

The mediation model tested whether the relationship between age and fluid intelligence could be (at least partially) accounted for by the relation between age and the brain response to difficult problem-solving. Mediation was assessed in Matlab using a set of linear regressions (Baron and Kenny, 1986; MacKinnon et al., 2007). The first equation below expresses the total linear relation between age and IQ; the second equation expresses the unique linear relation between age and IQ when also modelling the effect of the BOLD response on IQ:

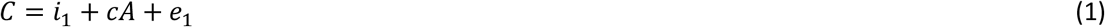

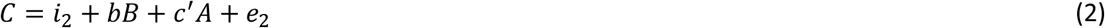

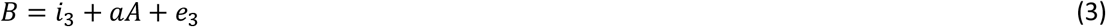

where the variable *A* is age, *B* is the brain’s BOLD response at a given voxel or ROI, and *C* is Culture Fair IQ; the coefficient *a* reflects the effect of age on the brain response, *b* reflects the effect of the brain response on IQ while controlling for age, *c* reflects the total effect of age on IQ, and *c’* reflects the ‘conditional direct’ effect of age on IQ while controlling for the brain response; *e*_*1-3*_ are residuals and *i*_*1-3*_ are intercept terms.

Where the data are consistent with mediation, *c’* would have reduced magnitude compared to *c*, i.e. the inclusion of the brain response in the model explains some of the variance in IQ that would otherwise have been explained by age. For a linear model, the difference between *c* and *c’* is equivalent to the product of *a* and *b* (MacKinnon et al., 2007), which describes the ‘indirect’ effect of age on IQ as the effect of age on the brain response combined with the (age-adjusted) effect of the brain response on IQ.

Mediation is traditionally tested and easiest to interpret when there is no interaction between the mediator and independent variable in predicting the outcome variable (as assumed in equation 2 above). Therefore, before testing for mediation we tested for ROIs or voxels where age and the fMRI response interact in predicting IQ, which would suggest that the brain response moderates (buffers or exacerbates) the direct effect of age on IQ. That is, we tested the interaction term (*d*) in the model:

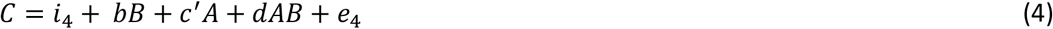

The significance of mediation can be assessed by separately testing the significance of *a* and the significance of *b*, or by testing the significance of the product of *a* and *b* directly (Baron and Kenny, 1986; MacKinnon et al., 2002). While the latter approach is more powerful under the null hypothesis that both *a* and *b* are zero, it has inflated type I error rates compared to a null hypothesis that either *a* or *b* might be zero (MacKinnon et al., 2002). We therefore used the more conservative conjunction of tests, which also allowed an efficient hierarchical approach in the context of testing multiple voxels across the brain. Specifically, we first identified voxels where the strength of the brain response (the potential mediator) showed a significant bivariate association with age (significant *a* coefficient in equation 3), thresholding for significance at p < 0.05, 2-tailed, while correcting for multiple comparisons (FDR). Of these voxels, we retained only those where there was also no evidence of moderation, defined as the interaction term (*d*) being both small (Cohen’s f^2^ < 0.02) and non-significant (p > 0.05, 2-tailed, without correction for multiple comparisons). This generated a conservative set of voxels within which classical mediation could then be tested based on additional significance of the *b* coefficient (equation 2). Since IQ has a *negative* relation with age, and we are interested in ‘consistent’ mediation (where the direct and indirect effects have the same sign, such that modelling the mediator reduces the size of the total effect; MacKinnon et al., 2000) we used a 1-tailed test that *b* had the opposite sign to *a* (i.e. IQ *increases* with brain response, which *decreases* with age, or vice-versa). Multiple comparisons were again accounted for by controlling the FDR <0.05, based on the conjunction of the tests for *a* and *b* (maximum p; Heller et al., 2007).

### Correlation of mediation effect size with strength of BOLD response to cognitive demand, across voxels

Testing whether voxels with the strongest mediation are also those most strongly activated by cognitive demands is complicated by spatial autocorrelation across the brain. Nearby voxels tend to have similar responses, so standard significance tests that assume independence of samples are invalid. We therefore tested significance using a Monte Carlo approach based on Moran eigenvector spectral randomisation (Wagner and Dray, 2015). The eigenvectors of a scaled proximity matrix derived from the Euclidian distance between voxels (Dray et al., 2006) comprehensively describe the spatial autocorrelation structure across all scales (Griffith and Peres-Neto, 2006). These eigenvectors were used to create a null model of the distribution of correlations that would be expected by chance, given the measured autocorrelation structure (Wagner and Dray, 2015). We used the Matlab implementation in the Brainspace toolbox (Vos de Wael et al., 2020), using the ‘singleton’ procedure and 10,000 random permutations.

### Diagnostic analyses of summary mediation model

For maximum sensitivity, diagnostic analyses of the mediation model were run using the mean BOLD signal across all voxels with significant mediation, data from all 252 participants, and a preferential response to difficult problem-solving. To confirm correct specification of the functional form of equations 1-3, i.e. that additional non-linear functions of the independent variables are not required to fit the data, we used the RESET test (Ramsey, 1969), as implemented in the Panel Data Toolbox (Alvarez et al., 2017). To test for heteroscedasticity of residuals from equations 1-3, i.e. whether residual variance varied as a function of the independent variables, we used the version of the White test proposed by Wooldridge (2012, p279).

Being based on regression between observed variables, relationships in the mediation model could potentially be induced indirectly by unmodelled variables that drive covariation between the observations. For example, if some participants were more distracted in the scanner this would likely lead to both lower performance and weaker task-induced BOLD signal. By using a measure of fluid intelligence acquired in a previous session, we avoid confounders that could affect simultaneous measures of performance and BOLD signal, and so ensure that the results generalise to a standard measure of fluid intelligence that is stable over time. Nonetheless, other potential confounds remain (e.g. general predisposition to distraction) and cannot be exhaustively excluded. Therefore, to assess the robustness of the mediation result to unmodelled confounders we performed a sensitivity analysis using the Left Out Variables Error (LOVE) method (Mauro, 1990; MacKinnon and Pirlott, 2015). This analysis asks how much the observed mediation strength might be overestimated in the presence of hypothetical confounders that correlate to varying degrees with the modelled variables. We expect age to be a cause rather than an outcome, and so to be unaffected by unmodelled variables. In this analysis, we therefore focused on potential confounding of the relationship between the brain and fluid intelligence measures. The analysis was implemented in Matlab based on the example in Valente et al. (2017).

The mediation analyses test whether the data are consistent with the hypothesised causal model (i.e. IQ and neural responsiveness are both affected by age, and IQ is also affected by neural responsiveness); however they cannot distinguish between alternative causal models (Fiedler et al., 2011). We therefore estimated the causal relationship between each pair of variables using a “linear non-Gaussian acyclic model” (LiNGAM), which can recover the causal directions under the additional assumptions of an acyclic model with no more than one error term being perfectly Gaussian (Shimizu et al., 2006). Briefly, the method starts from the observation that the matrix description of the linear relations between a set of centred random variables *X*:

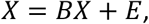

where *B* is a coefficient matrix and *E* are independent error terms, can be rearranged to:

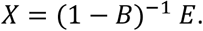

If *E* are assumed to be not perfectly Gaussian, then independent component analysis (ICA; Comon, 1994; Hyvarinen, 1999) can be used to decompose *X* into the independent error components multiplied by a mixing matrix, up to an undetermined scaling and permutation. The coefficient matrix *B* can then be derived from the mixing matrix, with the correct permutation and scaling determined from the assumptions that the model is linear and acyclic and so *B* should be lower-triangular. The model was estimated with an algorithm based on this ICA-plus-permutation approach, using the Matlab LiNGAM package (https://sites.google.com/view/sshimizu06/lingam). Since ICA-LiNGAM is not scale invariant, all variables were standardized to unit variance. The model was first estimated using the mean BOLD signal averaged across mediating voxels as defined above. Directionality of each path was defined as the estimated absolute connection strength in the hypothesised direction, minus the absolute connection strength in the reverse direction (of which only one is non-zero, given the acyclicity assumption). Thus positive values would reflect causality in the hypothesised direction. The reliability of this estimate was assessed using bootstrap (15,000 resamples). Since the bootstrapped sampling distribution was found to be biased with respect to the observed statistic, and far from Gaussian, it was not conducive to constructing confidence intervals. We therefore used the watershed algorithm to split the mediating voxels into 77 clusters (one per local minimum), ran LiNGAM on each cluster, discarded clusters where the sign of any undirected coefficient did not match that estimated from the all-voxel model (mean 44% across bootstrap resamples) or the LiNGAM algorithm warned that the coefficient matrix was not lower-triangular (mean 26% across bootstrap resamples), and calculated the mean directionality estimates across remaining clusters. This procedure was again repeated across 15,000 bootstrap resamples. In line with the central-limit theorem, using this mean estimate of directionality produced an approximately normal sampling distribution, which allowed a bias-corrected 95% confidence interval to be constructed.

### Moderated mediation analyses

We ran moderated mediation analyses to test whether the coefficients of the proposed mediation model depended on the level of a fourth variable (see Methods: “Lifestyle activities measures”). We used the approach of Edwards and Lambert (2007), allowing the moderator to affect any of the paths in the model, and testing for moderation of both single and compound paths. The analysis was implemented in Matlab, based on the example in Edwards and Lambert (2007). Simple effects were calculated for levels of the moderator one standard deviation above and below the mean. Differences between levels of the moderator were tested using standard parametric tests for the single paths (*a, b, c’*), and using bias-corrected percentile bootstrap (15,000 resamples) for the compound paths (indirect and total effects of age on fluid intelligence).

### Control analyses adjusting for head motion

Head motion is expected to increase with age and to decrease with fluid intelligence (Siegel et al., 2017), and is therefore a potential confound as well as a source of noise. While this is of particular concern for measures of functional connectivity (Ciric et al., 2018; Parkes et al., 2018) it can also degrade task fMRI data (Siegel et al., 2014). Therefore, on the suggestion of a reviewer, all analyses were repeated after regressing summary measures of individual differences in head motion. Two measures of head motion were calculated, using SPM Utility Plus (Pernet, 2021): the mean frame-wise displacement (FD) across the fMRI scan (Power et al., 2012) and the proportion of high-motion frames (FD > 0.9, with this threshold being suitable for task-based fMRI in cohorts with moderate motion; Siegel et al., 2014). All analyses were repeated after regressing these measures from the key variables of age, IQ and the BOLD response at each ROI/voxel, as well as lifestyle measures in analyses of moderated mediation. This carries a risk of removing actual effects of interest, to the extent that they happen to covary with head motion (Power et al., 2014); however, as a supplementary analysis it can identify results where head motion may be (or is unlikely to be) a confound, and could unmask results that might otherwise be obscured by the variance that head-motion shares with other variables.

## Results

### Behaviour

Fluid intelligence (IQ), measured using the Culture Fair test (Cattell and Cattell, 1973), had a mean of 106.6 and standard deviation of 19.5 (N=252; range 47 to 158 using the conversion in the manual, and 43 to 142 when estimated as a latent variable; see Methods). Since the test was originally constructed to have a population mean of 100 and standard deviation of 16, our higher mean may reflect the Flynn effect (Colom and Garcia-Lopez, 2003), while the higher variance may reflect our wide age range. It is unclear whether the lowest values reflect genuinely low fluid intelligence in older participants, or a qualitative difference in their ability to understand or perform the tasks. Since we had no principled reason to exclude the lowest performers, all participants were included in the analysis. However, on the suggestion of a reviewer, all analyses were repeated after excluding participants whose latent IQ score was more than three standard deviations below the mean (one participant in the ROI analysis plus a second in the subsequent analyses; manual-normed IQ scores 47 & 57; latent IQ scores 43 and 48; ages 71 & 86). The main conclusions were unchanged.

For the fMRI task, reaction time (RT) was calculated as the median time spent on each accurately answered problem at first attempt, per condition and participant. Participants were, as expected, substantially faster on the easy (mean RT = 1.02 s) than the difficult (mean RT = 4.83 s) problems (t(251)=37.9; p=7.4×10^−106^). Participants were also substantially more accurate on the easy (mean accuracy = 97.6%) compared to the difficult (mean accuracy = 56.2%) problems (t(251)=34.2; p=2.4×10^−96^).

### fMRI difficulty contrast

The group mean BOLD response associated with difficult versus easy problem-solving is shown in Figure 1B. As expected, the difficult condition produced greater activation of the MDN bilaterally, extending into occipital cortex which may reflect enhanced attention to the visual stimuli. The difficult condition was also associated with reduced response of the DMN, auditory, and sensorimotor cortex. Reduced responses in sensorimotor cortex, especially in the left hemisphere, are expected due to the less frequent (right-hand) button presses; reduced activation in auditory cortex may reflect attentional suppression of the scanner noise with increased focus on the visual modality.

### Bivariate relationships between age, fMRI, and fluid intelligence

We start by describing the three bivariate relationships in the full sample. Firstly, the association between age and fluid intelligence (Figure 2, bottom) showed strong and approximately linear cross-sectional decline across the age range, as reported previously using related measures and overlapping samples of Cam-CAN participants (Kievit et al., 2014; Samu et al., 2017). The Pearson correlation coefficient (r = -0.66) corresponded to an average loss of 7.2 IQ points per decade of age.

**Figure 2:**
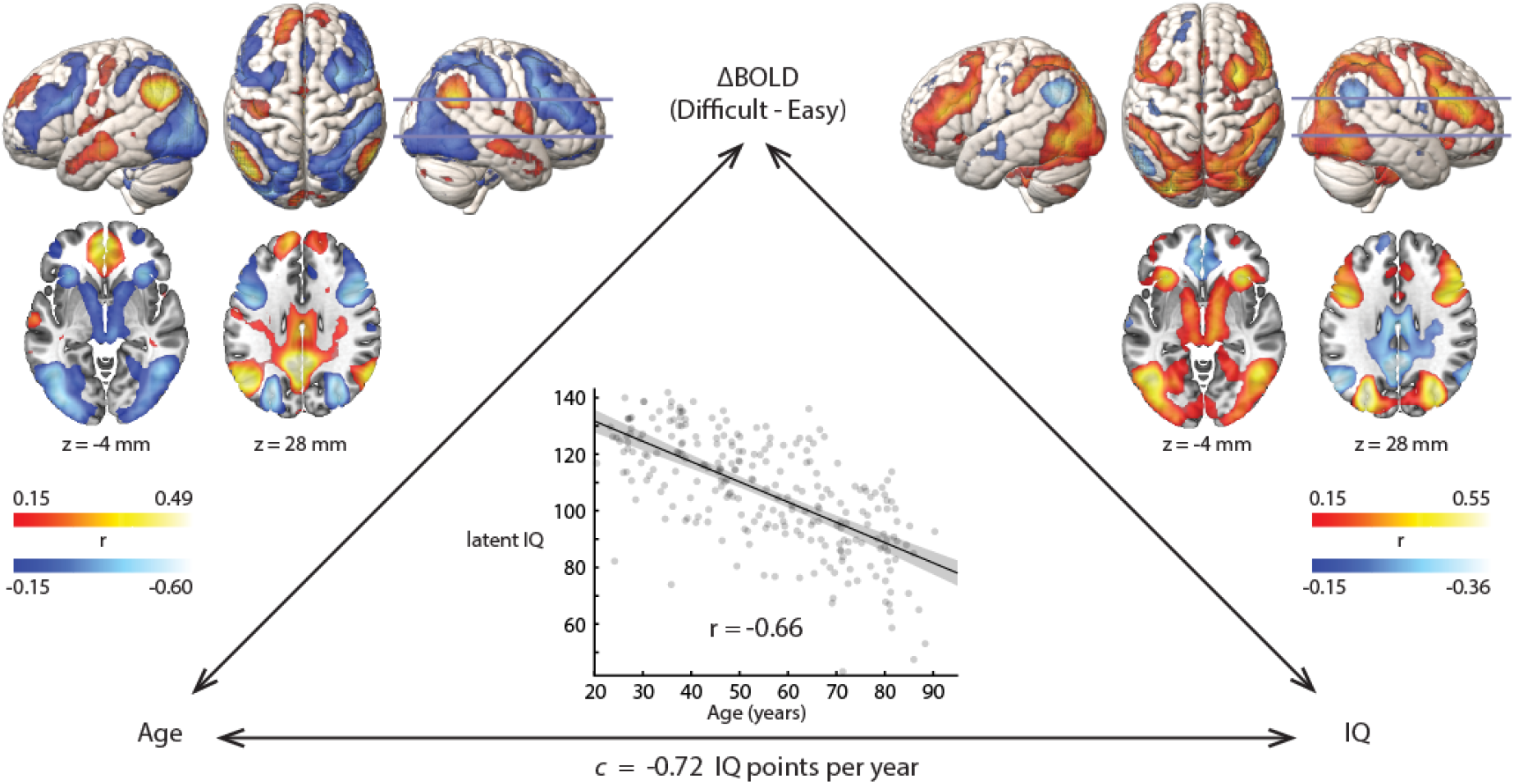
Bivariate relationships between age, fMRI response, and fluid intelligence (N=252). Voxel-wise correlation coefficients are thresholded based on significance at FDR < 0.05. Slice positions are labelled in MNI coordinates and marked on the right hemisphere rendering. Surface renderings show activations to a depth of 15 mm.

Secondly, we assessed how the fMRI response to difficult problem-solving depended on the age of the participants (Figure 2, left). To a first approximation, the results corresponded to a weakening of the typical response, as age increased: regions that were activated by the difficult condition on average (warm colours in Figure 1B) showed less activation (cool colours in Figure 2, left), while DMN regions that were typically suppressed in the difficult condition showed less suppression. We note two partial exceptions to this general pattern: auditory and sensorimotor cortex, which were strongly suppressed in the difficult condition, showed relatively less difference across age, or even increased suppression (anterior IPS); similarly, parts of the basal ganglia showed increasingly negative response to difficulty with age, despite the difficulty response being non-significant (caudate) or weakly negative (putamen) on average.

Thirdly, we assessed how the fMRI response to difficult problem-solving correlated with individual differences in fluid intelligence (Figure 2, right). The pattern was remarkably similar to the correlation with age, but inverted in sign. That is, regions whose activity was associated with higher fluid intelligence tended to be those with the greatest age-related decline in activity, and vice versa. One exception was again auditory and sensorimotor regions: although these were suppressed in the difficult condition, and suppression somewhat reduced with age, we saw little association with IQ, consistent with the expectation that these regions are responding to specific sensory and motor aspects of the particular task design, rather than having a more general role in fluid intelligence.

### Mediation of age-related differences in IQ by neural response to cognitive demand: Network-based replication

Given the pattern of bivariate relationships, it is possible that, for some voxels, age independently impacts fluid intelligence and the BOLD response. To identify regions where a reduced response to difficult problem-solving potentially mediates the effect of age on fluid intelligence, we jointly predict IQ from both age and the fMRI response, and test for a decrease in the remaining (direct) effect of age on IQ, relative to the total effect of age on IQ when modelled alone. In other words, we identify regions where variation in IQ is partly explained by an indirect path of age affecting the brain which has a consequent effect on IQ. This also entails the relationship between the fMRI response and IQ remaining significant when controlling for age.

We first sought to replicate the recent finding that the responsiveness of frontoparietal regions is consistent with such mediation of age differences in fluid intelligence (Samu et al., 2017). For this analysis we thus restricted ourselves to the independent subset of participants who were not included in the preceding study (N=154), and we summarised the brain response according to major cortical networks (Ji et al., 2019) plus more focussed definitions of core multiple-demand (Assem et al., 2020a) and default mode networks. An initial test for an interaction between age and brain response in predicting IQ found no significant effects for any network after adjusting for multiple comparisons (all FDR-adjusted p > 0.64). Therefore, consistent with Samu et al. (2017), there was no evidence that the relationship between the neural response and fluid intelligence differed with age.

We then probed neural mediation of the age effect on fluid intelligence. Significant mediation was observed in four networks: Dorsal-attention, Secondary-visual, Frontoparietal, and the Core MDN (which is a subset of the Frontoparietal network), confirmed by joint significance of their *a* and *b* paths (Table 1) as well as bootstrapped confidence intervals around their product (Figure 3A). We therefore replicate the mediation observed by Samu et al. (2017), link it to particular functional networks, and extend it from mediation of concurrent task performance to mediation of participants’ fluid intelligence more generally. The mediation effect size (*ab*) is plotted against the group-average fMRI response to difficult versus easy problem-solving in Figure 3A. The same data are plotted a different way in Figure 3B, breaking down each mediation effect into the magnitude of the *a* and *b* coefficients. This shows that the core DMN is impacted by age (*a*) at least as strongly as the networks activated by the difficult condition (red); the reason that it does not significantly mediate the decline in IQ is due to its small effect on IQ after controlling for age (*b*), again replicating Samu et al. (2017). Overall, it is striking that evidence for mediation is specific to those networks responding most positively to task difficulty. This is consistent with correlation of the BOLD response with the *a* and *b* paths separately, reported across ICA components by Samu et al. (2017).

**Table 1.**
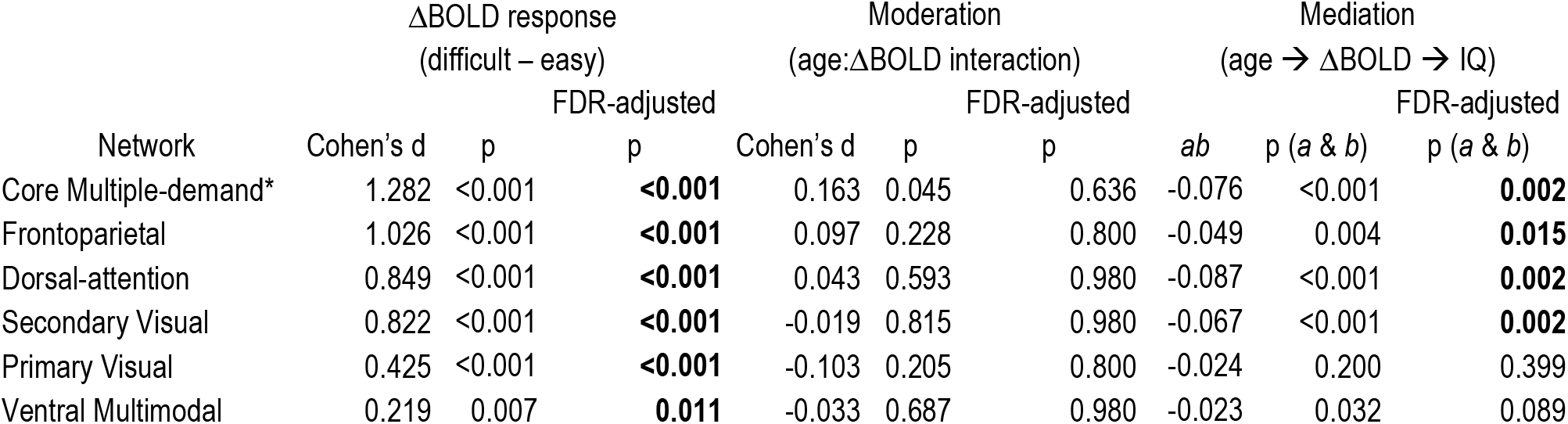
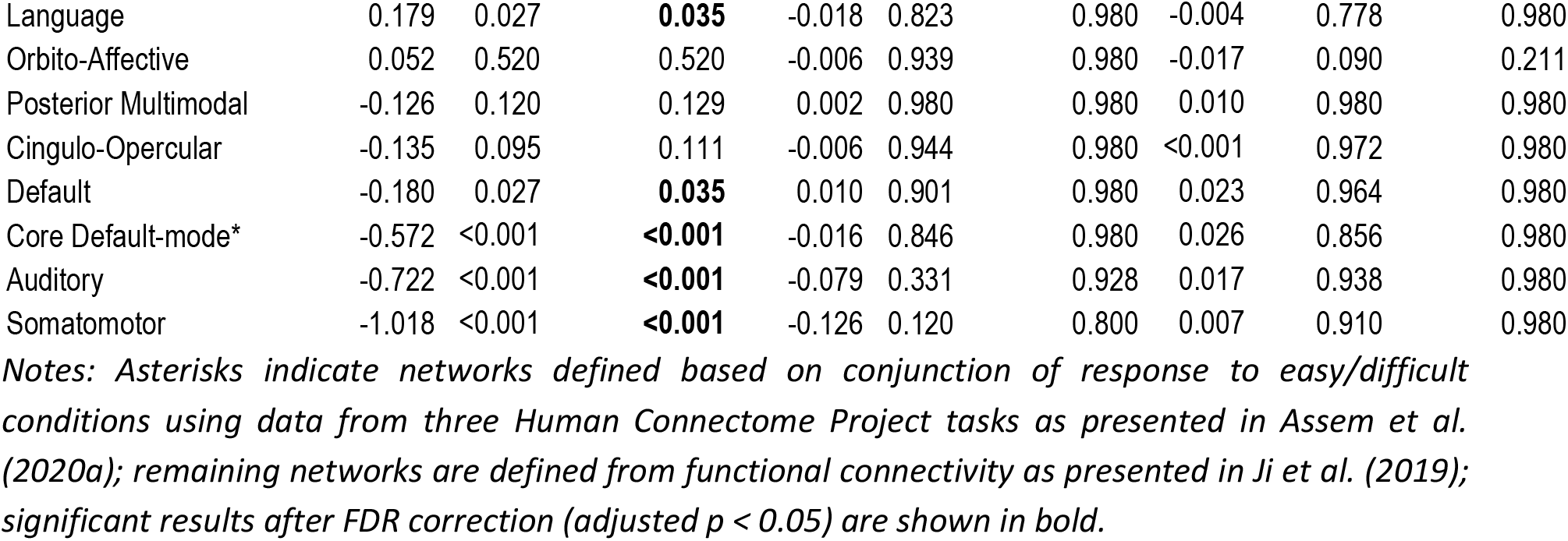
BOLD response to cognitive difficulty, and its moderation and mediation of age effects on fluid intelligence, for cortical network ROIs (N=154).

**Figure 3.**
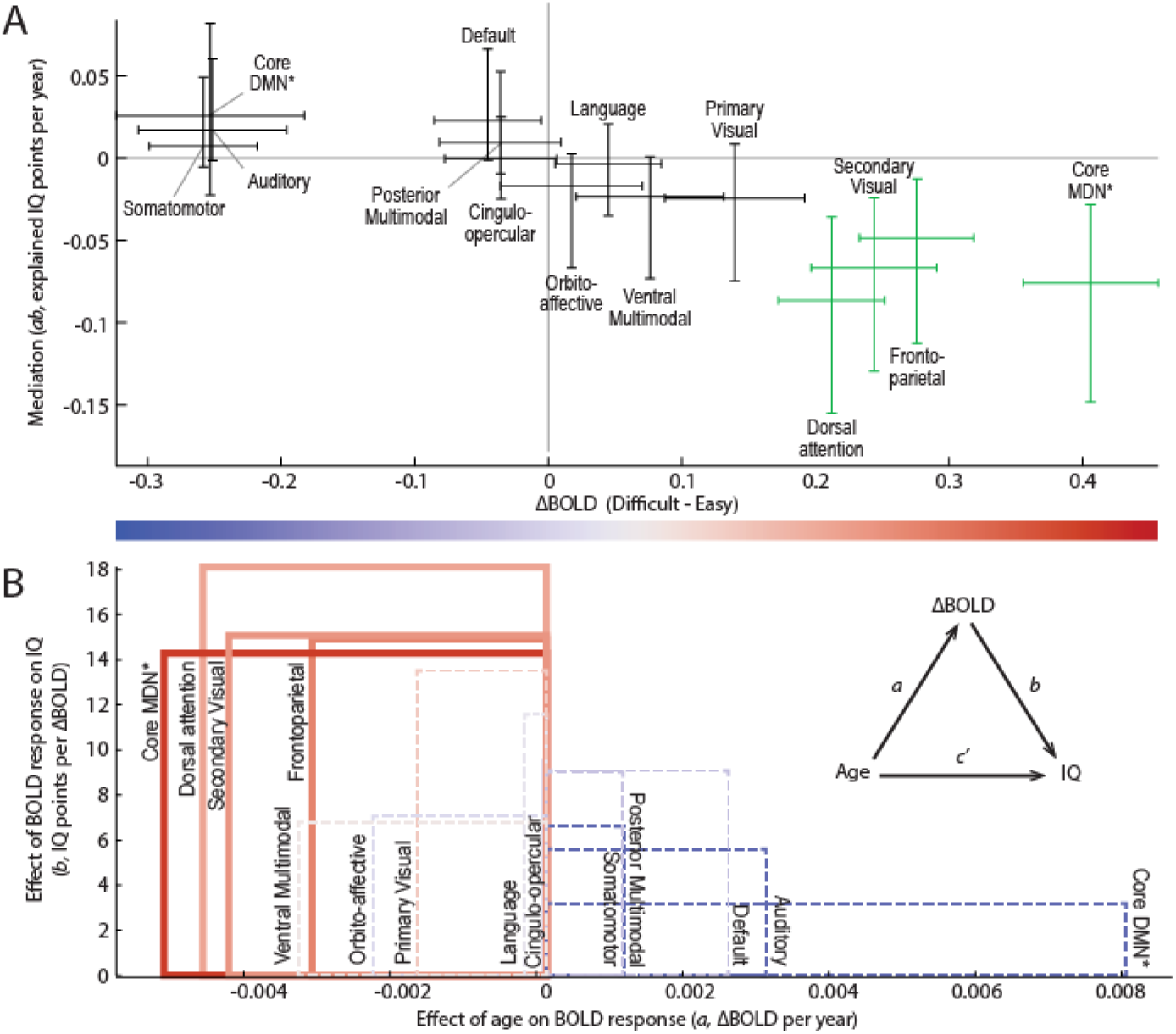
BOLD response to cognitive difficulty, and its mediation of age effects on fluid intelligence, for cortical network ROIs (N=154). A) Overall mediation effect size versus group average BOLD response. Error bars are 95% confidence intervals derived from the associated t-test (BOLD contrast) or bias-corrected bootstrap (mediation). Networks with significant mediation after FDR correction (adjusted p < 0.05) are shown in green. B) The b coefficient (reflecting the age-adjusted effect of the BOLD response on IQ) is plotted versus the a coefficient (reflecting the effect of age on the BOLD response), with group-mean BOLD response indicated by colour. The inset depicts the mediation model. The area of each rectangle conveys the mediation effect size (ab) for the corresponding network. Solid and dashed lines indicate significant and non-significant mediation, respectively. Asterisks indicate networks defined from their conjunction response to easy/difficult conditions using data from three Human Connectome Project tasks as presented in Assem et al. (2020a); remaining networks are defined from functional connectivity as presented in Ji et al. (2019).

The mediation results considered so far are at a relatively coarse spatial scale, employing combined ICA-based brain-wide spatial components (Samu et al., 2017) or functional/anatomical networks (our replication). Next, we combine both samples to increase power and precisely localise the mediation effect on a voxel-wise basis. This also allows us to quantify the degree to which mediation is associated with voxels that respond positively to task difficulty, separately for voxels activated and voxels suppressed by cognitive demand, and accounting for spatial autocorrelation.

### Voxel-wise localisation of mediation effect, and its association with voxels most responsive to cognitive demand

To identify mediating voxels in a conservative but efficient manner, we took a hierarchical approach where we first tested for significance of the *a* path, and used significant voxels (after FDR correction) to define an analysis mask within which to test for voxels where the *b* path was also significant. We further restricted the analysis mask to voxels where there was no evidence of an interaction between age and brain response in predicting IQ. Consistent with Samu et al. (2017) and with our independent replication at the network level, no voxels showed such an interaction after correcting for multiple comparisons (all FDR-adjusted p > 0.32). Nonetheless, to be conservative we excluded voxels based on uncorrected significance (p < 0.05) or more than ‘small’ effect size (Cohen’s f^2^ > 0.02) of the interaction term.

The resultant analysis mask is shown as the blue and red overlays in the second row of Figure 4A, where the colour indicates the sign of the difficulty contrast (red for difficult > easy; blue for easy > difficult). Voxels with significant mediation after correcting for multiple comparisons within this mask (FDR < 0.05) are shown in green. Interestingly, over 90% of voxels with significant mediation responded more to difficult than to easy problem-solving. Identified voxels show a close relationship to MDN regions, including key foci along the precentral sulcus and middle frontal gyrus, and within the intraparietal sulcus, anterior insula, and anterior cingulate cortex. Mediating voxels are also observed in lateral occipital cortex and subcortical structures including the basal ganglia, thalamus and cerebellum, which are often co-activated with the cortical MDN (Assem et al., 2020a).

**Figure 4:**
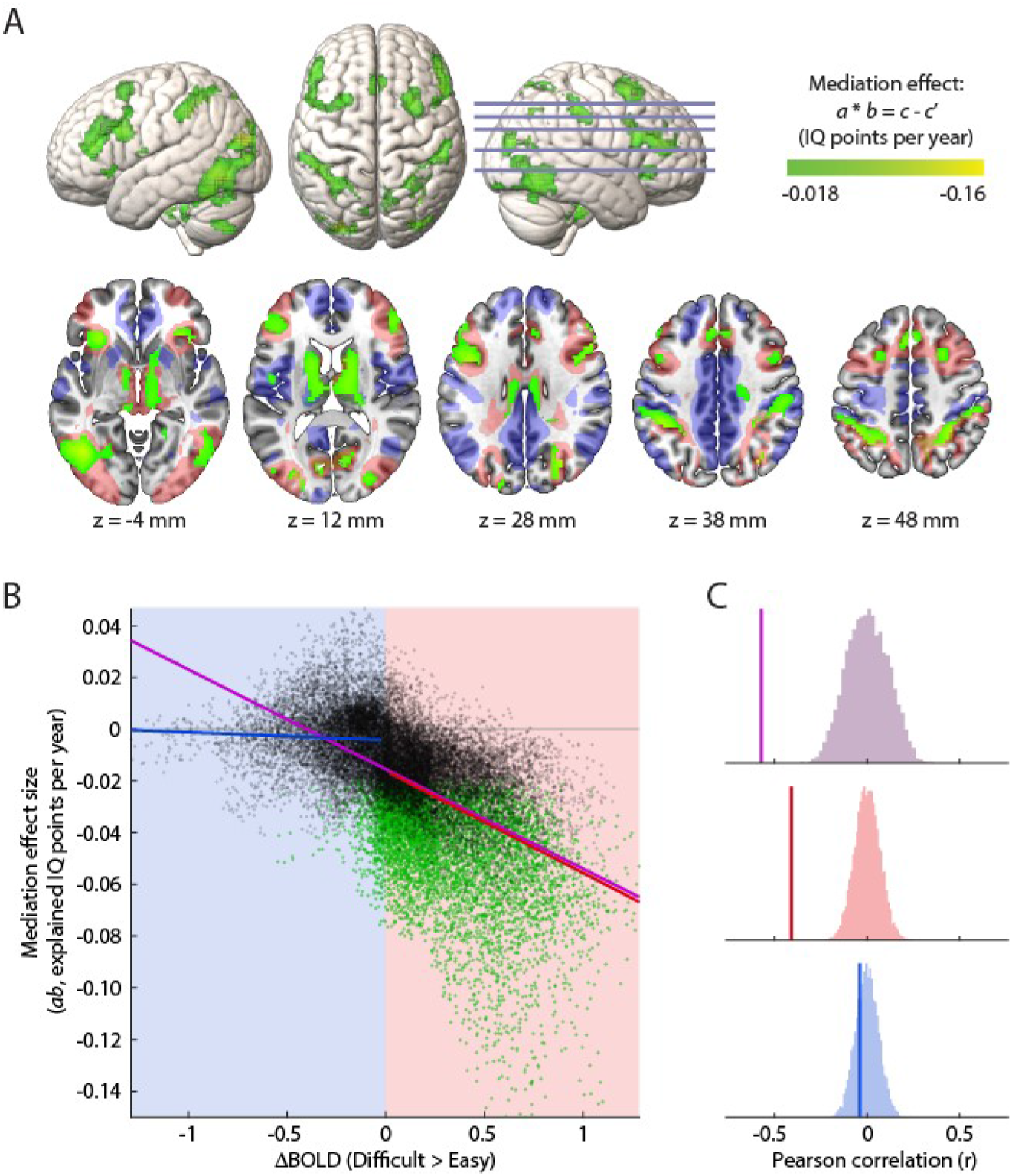
Voxel-wise mediation analyses. A) Regions with significant mediation (FDR < 0.05 within analysis mask) are shown in green, overlaid on surface renderings (top) and horizontal slices (bottom). Slice positions are labelled in MNI coordinates and marked on the right hemisphere rendering. In the bottom row, the analysis mask is shown by the blue/red overlay, coloured by the sign of the difficulty contrast (red for difficult > easy; blue for easy > difficult). Surface renderings show activations to a depth of 30 mm. B) Scatterplot showing the relationship between the size of the mediation effect and the group-mean difficulty contrast. Each point is a voxel within the analysis mask, coloured green where mediation is significant (FDR < 0.05). C) Pearson correlations (vertical lines) between the size of the mediation effect and the difficulty contrast, compared to permutation null distributions constructed using Moran spectral randomisation. The correlation is separately tested across all voxels (purple), only voxels with a positive response to task difficulty (red), and only voxels with a negative response to task difficulty (blue), corresponding to the lines of best fit in panel B.

The relationship between neural mediation of fluid intelligence and a voxel’s average response to difficult problem-solving is illustrated in Figure 4B, where the mediation effect size is plotted against the response to difficult vs easy problems. The overall relationship across voxels is highly significant (Pearson correlation, r = -0.57; p < 0.001; Figure 4B-C, purple) after using a permutation test with Moran spectral randomisation to comprehensively account for spatial autocorrelation in the maps (Wagner and Dray, 2015; Vos de Wael et al., 2020). We also test the correlation separately within voxels significantly activated by difficult versus easy problem-solving, and within voxels significantly suppressed by difficult versus easy problem-solving. The correlation is again significant within positively activated voxels (r = -0.41; p < 0.001; Figure 4B-C, red) whereas there is no evidence that mediation is correlated with the level of difficulty-induced suppression (r = -0.04; p = 0.52; Figure 4B-C, blue).

### A summary mediation model, and diagnostic analyses: robustness to confounders and a test of causal direction

To summarise and illustrate the localised mediation effect, we averaged across those voxels with significant mediation, data from all 252 participants, and a preferential response to difficult problem-solving. For these voxels, the mediation model is depicted in Figure 5A. Figure 5B illustrates the mediation effect in terms of the underlying relationships as suggested by MacKinnon et al. (2007), where the green triangles highlight the reduction of the total effect of age when modelling the neural response (c – c’).

**Figure 5:**
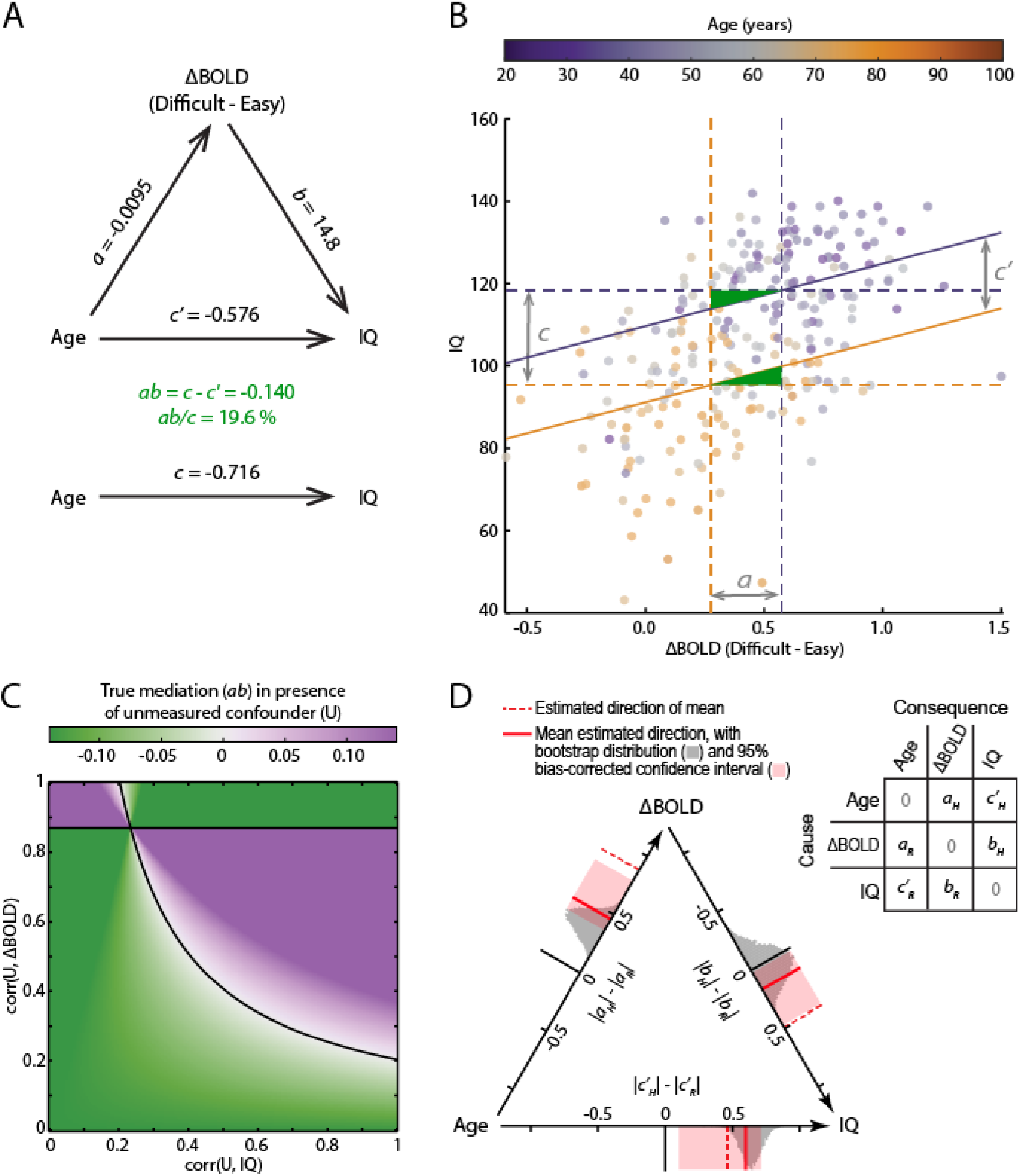
Summary and assessment of the localised mediation effect, averaged across voxels where mediation is significant, data are available from all participants (N=252), and the difficult-easy contrast is positive. A) Fitted mediation model. B) Illustration of the mediation effect and underlying relationships. Vertical dashed lines represent equation 3 at the first and third quartiles of the age distribution (purple=39 years; orange=71 years); horizontal dashed lines represent equation 1 at the same ages; solid lines represent equation 2 at the same ages, with their slope equal to coefficient b; grey arrows reflect coefficients a, c and c’, when age is scaled by the interquartile range; the standardized mediation effect size is indicated by the green triangles, relative to the area of the central rectangle, which can be seen to account for about a fifth of the total effect. C) Left Out Variables Error (LOVE) plot indicating how large any correlations with an unmodelled confounder (U) would need to be to reduce the true mediation to zero. The observed mediation effect size lies at the origin. D) Estimated causal direction of each path, using ICA-LiNGAM after standardization of variables (see Methods). Axes represent the difference of absolute model coefficients for the hypothesized (H) minus reverse (R) causal directions, indicated by the matrix. Estimation of the model returns three non-zero coefficients in this matrix, one for each pair of variables, in either of the two possible directions. (Zeros on the diagonal reflect the fact that variables are assumed not to cause themselves.) Dashed red lines were calculated using the mean signal across all mediating voxels as plotted in B; solid red lines represent mean direction estimates, averaged across local clusters of mediating voxels whose undirected coefficient signs matched those from the mean signal; grey histograms show the bootstrap sampling distribution of the cluster-mean directionality; pink bands show bootstrapped 95% bias-corrected percentile confidence intervals for the cluster-mean directionality.

Since the mediation effect rests on standard assumptions of linear regression, we next confirmed that the model is appropriate. We used the RESET test (Ramsey, 1969) to confirm correct specification of functional form, i.e. that non-linear functions of the independent variables were not required to fit the data. For equations 1-3 we found no evidence of misspecification (all F(3,246-7) < 2.23, all p > 0.08). Although the linearity of relationships with age may break down at the extremes of the lifespan, approximate linearity is common for accuracy-based performance measures (Salthouse, 2011a) and BOLD responses (Grady et al., 2006) during healthy aging. Next we used the White test (Wooldridge, 2012) to assess whether the residual variance varied as a function of the independent variables. For equations 1-3 we found no evidence against homoscedasticity (all *χ*_2_(2) < 4.36, all p > 0.11). We also note that any violation of the assumption that variables are measured without error would underestimate the magnitude of the true mediation effect (Pieters, 2017).

The mediation analysis assumes that there are no confounding variables omitted from the model (MacKinnon and Pirlott, 2015). Potential unmeasured confounders, however, are plausible, if not inevitable, in practice. Assuming that chronological age must be a cause rather than a consequence, the main concern is that an unmodelled variable could produce a spurious mediation effect by covarying with both IQ and the neural measure (Salthouse, 2011a). Using a trait measure of fluid intelligence rather than concurrent performance avoids some potential sources of shared variance, but others will remain. Therefore, to infer a direct relationship between neural responsiveness and fluid intelligence, and to establish the validity of the mediation model as a potential mechanistic explanation, it is important that the mediation result is robust to possible unmodelled covariates. To address this, we used sensitivity analysis (Mauro, 1990; Valente et al., 2017) to estimate how strong any unmodelled confounders would need to be to fully explain the observed mediation relationship. Figure 5C plots the size of the true mediation effect as a function of an unknown confounder’s correlation with the fMRI response and with IQ. This shows that for there to be no true mediation (at points along the black line) the correlations of any confounder with both the fMRI response and IQ would need to exceed 0.43 on average (and at least one must exceed 0.45). While this cannot be ruled out, it provides reassurance that the mediation result is robust to the possibility of moderate confounding.

Although the mediation analyses confirm that the data are consistent with the hypothesised causal model, without experimental manipulation of the variables the data are equally consistent with alternative causal models (Fiedler et al., 2011; Salthouse, 2011a; MacKinnon and Pirlott, 2015). The various possible causal orderings are statistically equivalent, having the same covariance matrix and global model fit, and one cannot adjudicate between them based on the size or significance of their mediation effect (Thoemmes, 2015). To assess causal directionality, we therefore used a “linear non-Gaussian acyclic model” (LiNGAM; Shimizu et al., 2006). By making additional assumptions that no more than one error term is perfectly Gaussian, and that the model is acyclic, LiNGAM uses ICA to estimate the generating causal model based on purely observational data (see Methods). Using the mean across mediating voxels, as defined above, LiNGAM estimated the causal directions to be in line with the assumed model (Figure 5D, dashed red lines). Bootstrap resampling, however, revealed the sampling distribution to be biased and far from multivariate normality. We therefore split the mediating voxels into local clusters, applied LiNGAM to each cluster, discarded clusters where a causal model could not be reliably identified or whose undirected model coefficients differed in sign from the all-voxel model, and averaged directionality estimates across the remaining clusters. Again, estimated causal directions were all in line with the assumed model (Figure 5D, solid red lines), now with approximately Gaussian bootstrapped sampling distributions (grey histograms) from which 95% bias-corrected percentile confidence intervals were constructed (pink bands). Note that the bootstrap distributions remain shifted with respect to their observed direction estimates (solid red lines), suggesting that the observed estimates are similarly biased with respect to their true population values; therefore the bias-corrected confidence intervals do not lie on the percentiles of the sampling distribution but are shifted in the opposite direction to compensate (Hesterberg, 2015). None of the intervals span zero, suggesting confidence in the estimated directions under the assumptions of the model, although the interval for the brain-behaviour relation is close to zero and the assumptions in this case are questionable (see Discussion).

### Moderation of the mediation model by a varied active lifestyle

Finally, in exploratory analyses, we tested whether any path in the observed mediation model might be moderated by self-reported regular physical activities, in terms of their variety (number of different activities; range 0-14), mean frequency per activity (episodes per year; range 12-365), mean duration per episode (range 4-415 minutes), or total duration (range 10-3240 hours per year). We found that the variety of physical activities significantly moderated the path from the brain response to IQ, but not the effect of age on the brain response, or the conditional direct effect of age on IQ (Table 2). This carried through to a significant moderation of the overall indirect effect of age on IQ, although moderation of the total effect of age was not significant. In contrast to the moderating effect of the variety of activities, the mean frequency and duration of each activity had no significant effect on any path in the model, nor did the total duration of activity.

**Table 2.** Moderated mediation results. Moderation of each single and compound path in the mediation model, by various moderator variables.

| Moderator | N | 1<sup>st</sup> stage of mediation path |  | 2<sup>nd</sup> stage of mediation path |  | Direct effect |  | Indirect effect (mediation) |  | Total effect |  |
| --- | --- | --- | --- | --- | --- | --- | --- | --- | --- | --- | --- |
| | | $\Delta a$ | p | $\Delta b$ | p | $\Delta c'$ | p | $\Delta(ab)$ | p | $\Delta(ab+c')$ | p |
| Physical activities: |  |  |  |  |  |  |  |  |  |  |  |
| Variety | 239 | -0.000046 | 0.983 | -17.1 | **0.0061** | 0.0077 | 0.941 | 0.157 | **0.0095** | 0.165 | 0.146 |
| Frequency per activity | 231 | -0.00033 | 0.876 | 4.67 | 0.499 | -0.081 | 0.501 | -0.048 | 0.495 | -0.128 | 0.206 |
| Duration per episode | 220 | -0.0028 | 0.276 | -7.52 | 0.189 | -0.015 | 0.921 | 0.037 | 0.587 | 0.023 | 0.888 |
| Total duration | 220 | 0.00035 | 0.854 | -7.30 | 0.229 | -0.745 | 0.424 | 0.074 | 0.309 | -0.00072 | 0.811 |
| Variety* | 220 | -0.00093 | 0.701 | -15.7 | **0.022** | 0.134 | 0.244 | 0.138 | 0.076 | 0.272 | **0.011** |
| Frequency per activity* | 220 | -0.0013 | 0.531 | -4.88 | 0.502 | -0.020 | 0.857 | 0.031 | 0.678 | 0.010 | 0.930 |
| Duration per episode* | 220 | -0.0032 | 0.211 | -10.1 | 0.096 | 0.089 | 0.519 | 0.057 | 0.522 | 0.146 | 0.295 |
| Total duration* | 220 | 0.0013 | 0.499 | 5.42 | 0.373 | -0.087 | 0.348 | -0.036 | 0.586 | -0.123 | 0.342 |
| Non-physical activities: |  |  |  |  |  |  |  |  |  |  |  |
| Variety | 237 | -0.0040 | 0.101 | -11.5 | 0.074 | -0.087 | 0.500 | 0.049 | 0.468 | -0.038 | 0.769 |

Notes: \*Asterisks indicate moderators that are residualised with respect to each other. $\Delta$ indicates the change in coefficients across two standard deviations of the moderator variable. Following Edwards and Lambert (2007), p-values for single paths are based on standard errors from the regression model; p-values for compound paths (indirect and total effect) are based on bias-corrected confidence intervals derived from bootstrap resampling. Significant results ( $p < 0.05$ ) are shown in bold. Sample size (N) varies because not all participants answered all activities questions.

Some dependence between the four activity measures is expected. We therefore repeated the analyses using the residuals for each measure after regressing out the other three. The results were largely similar, again with an effect of variety, but no effects of frequency, mean duration or total duration. Now the total effect of age on IQ was also significantly moderated by the variety of activities, driven, as before, by reduction of the effect of brain response on IQ, although moderation of the compound indirect path no longer reached significance. The moderating effect of the variety of regular physical activities is illustrated in Figure 6, showing the change in simple slopes for each single and compound path (Edwards and Lambert, 2007). The substantial mediation effect for people engaging in relatively few regular physical activities (solid lines) is abolished for people engaging in a larger variety of activities (dashed lines), due primarily to decoupling of the relation between the neural response to task difficulty and fluid intelligence (blue lines).

**Figure 6:**
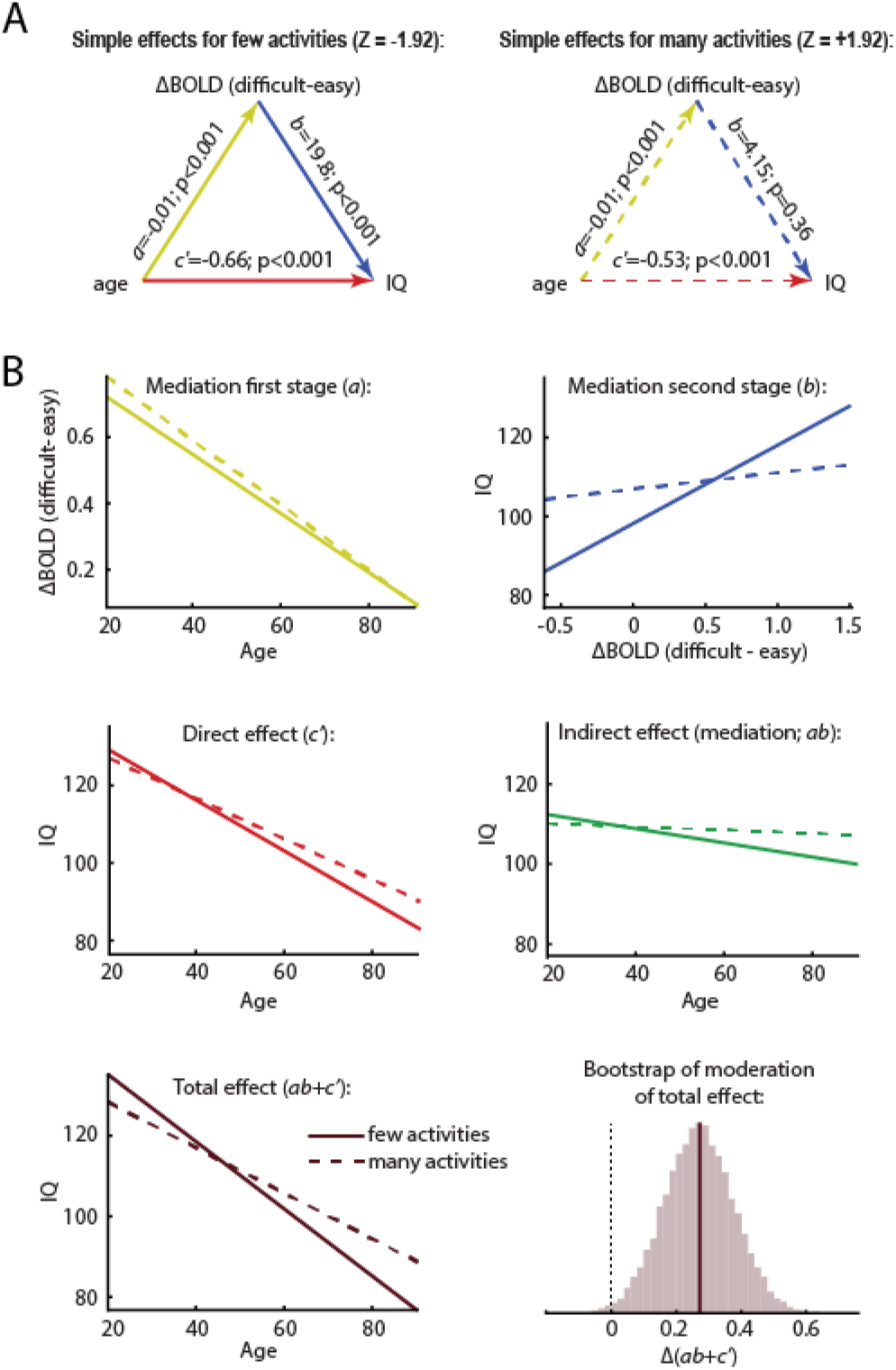
Moderated mediation. The moderator variable (Z) is the variety of regular physical activities, residualised with respect to the other physical activity measures. A) Simple mediation effects, at low (left, solid lines) and high (right, dashed lines) levels of the moderator (+/-1 standard deviation from mean). B) The effect of the moderator on each individual and compound path in the model. The solid and dashed lines again indicate low and high levels of the moderator respectively, as in panel A, with corresponding colours for the single paths. The final histogram shows the distribution of bootstrap samples used to calculate the p-value for the moderation of the overall effect of age on IQ.

In addition to physical exercise, engagement in socially and intellectually stimulating leisure activities is also thought to be beneficial for cognitive aging (Hughes et al., 2010; Kuiper et al., 2015; Yates et al., 2016; Borgeest et al., 2020). Therefore, given the significant moderating effect of the variety of regular physical activities, we next asked whether this generalises to the variety of regular non-physical activities, again measured by questionnaire, as well as in a home interview (number of social or intellectual activities; range 4-43). This time there was no moderating effect on any path in the model (Table 2).

### Control analyses adjusting for head motion

Individual differences in head motion can covary with both age and fluid intelligence (Siegel et al., 2017), and degrade task fMRI data (Siegel et al., 2014). Therefore, the potential contribution of head motion was assessed by repeating all analyses after regressing summary measures of mean framewise displacement (FD) across the fMRI scan, and the proportion of high-motion frames. These measures were highly related (r=0.91) and they both correlated substantially with age (r > 0.42, p < 1.4×10^−12^) and IQ (r < -0.39, p < 5.3×10^−13^). After regressing both motion measures from all key variables (age, IQ, BOLD response per ROI/voxel, and lifestyle measures), conclusions were largely unchanged, except as follows. In the ROI analysis, the mediation effect in the Frontoparietal Network ROI became non-significant, though the effect in the Core Multiple-Demand network remained. The percentage of significantly mediating voxels that were more active in the difficult condition increased from 90% to 95%. The size of the voxel-summary mediation effect increased from 19.6 % to 22.8 % of the total effect. In the LiNGAM analysis, estimated mean causal directions remained in the hypothesized direction, although the age∼IQ directionality no longer differed significantly from zero.

The largest differences were in the analysis of moderation by everyday activities. Here, the variety of physical activities continued to significantly moderate the second stage of the mediation path (p=0.010 when modelled alone; p=0.033 after covarying the other measures), but now alongside additional moderation by the residual duration of activity per episode (p=0.043), and without significant moderation of the total effect (p=0.137). Most interestingly, the first stage of the mediation path was now significantly moderated by the number of non-physical activities (p=0.008). This latter effect reflected an exacerbated age effect for people reporting more activities: the BOLD response to difficult problems increased with the number of activities for younger people (less than about 50 years old) whereas it decreased for older people who reported more activities. This effect of non-physical activities therefore differed from the effect of physical activities in two ways – it was associated with a different stage of the mediation pathway, and instead of flattening the age effect (Fig 8, blue lines), it steepened it (see https://osf.io/xgw56/ for results tables from these analyses).

In summary, the key conclusions were unchanged, suggesting that head-motion is unlikely to explain the mediation of age-related IQ decline by MDN responsiveness, the association of mediation with voxels responding positively to cognitive challenge, or its moderation by the variety of physical recreation. The emergence of additional lifestyle moderators in this control analysis suggests that head motion may act as a suppressor variable in these cases (MacKinnon et al., 2000), being associated with multiple other variables in opposite directions that can cancel out if not statistically adjusted. This suggests, in turn, that future studies would benefit from careful control of head motion, and that to further understand the complicated relationships that underlie the moderated mediation results it would be useful to replicate them, ideally via direct experimental manipulation.

## Discussion

This paper presents five key results. Firstly, we provide an independent replication that responsiveness of multiple-demand cortex to cognitive demand partially mediates age-related decline in fluid intelligence. Secondly, strongest mediation is specific to voxels most activated by cognitive demand, and not those suppressed by cognitive demand. Thirdly, the summarized mediation effect is robust to moderate confounding by unmodelled variables. Fourthly, assuming unidirectional causality, differences in brain response more likely drive IQ differences than vice-versa. Finally, diversity of physical activity moderates the summarized mediation effect, through decoupling of IQ from neural responsiveness to cognitive demand.

The specificity of mediation to the MDN and associated regions is notable, supporting previous hypotheses that “deterioration in DLPFC may (at least partially) underlie the relationship between adult age and abstract reasoning ability” (Phillips and Della Sala, 1998). However, MDN and DMN responses often anti-correlate (Fox et al., 2005), and this anti-correlation can positively or negatively relate to intelligence (Hearne et al., 2016; Santarnecchi et al., 2017b). Task-induced DMN suppression also reduces with age (Grady et al., 2006; Turner and Spreng, 2015), as does its coupling with frontoparietal attention networks (Spreng et al., 2016), while maintenance of DMN deactivation may explain cross-domain differences in cognitive aging (Samu et al., 2017). Although DMN suppression in this task indeed reduced with age, we found, matching Samu et al. (2017), no evidence that this mediated age-related IQ differences. With mediation strength selectively associated with voxels most activated, but not those most suppressed, by cognitive demand, reduced MDN function appears more important than altered DMN function in explaining age-related differences in IQ, while links between DMN activity and IQ may reflect confounding variables such as age. It remains possible that altered DMN activity may mediate age-related differences in other cognitive domains for which this network is specialised.

We observed no reliable indication that age moderates the association between neural responsiveness and IQ. This is perhaps surprising given the wide age range examined and the expected plasticity of neural recruitment and cognitive strategy (e.g. Cabeza, 2002; Davis et al., 2008; Park and Reuter-Lorenz, 2009), but matches previous observations using this task (Samu et al., 2017). Plausibly, neurocognitive shifts might primarily occur in more domain-specific tasks, when recruitment of domain-general MDN regions is not essential, but offers an optional compensatory strategy. Conversely, fluid intelligence may be especially susceptible to normal aging due to limited capacity for functional plasticity of the networks involved.

Multiple mechanisms undoubtedly link aging and fluid intelligence (Kievit et al., 2016). Indeed, the observed partial mediation explains around 20% of the relationship between age and IQ, leaving much room for additional mechanisms, alongside neural responsiveness to cognitive demand. For example, various aspects of structural brain integrity may play mediating roles (Salthouse, 2011a; Kievit et al., 2014). Network connectivity is also associated with fluid intelligence (Cole et al., 2012; Barbey, 2018; Dubois et al., 2018; Hilger et al., 2020) and changes with age (Tsvetanov et al., 2016; Bethlehem et al., 2020). Lifespan differences in cerebral vascularisation may further influence neural function and neurovascular coupling (West et al., 2019; Tsvetanov et al., 2021), although blood flow variation explains relatively little covariance between age, BOLD response and performance on this task (Wu et al., 2021). Future work could usefully address the relative importance of multiple mediators, and the relationships between them (Hedden et al., 2016).

While mediation analysis can suggest a mechanistic explanation for an observed relationship, it tests consistency with a hypothesised causal model, rather than inferring the causal structure itself (Fiedler et al., 2011; MacKinnon and Pirlott, 2015; Thoemmes, 2015). Although the LiNGAM method estimated directionality matching the hypothesised model, its assumption of non-reciprocal influences is tenuous in practice, and significance was sensitive to control analyses. Longitudinal data would provide another means to strengthen causal interpretations; if correlated changes are temporally separated, the preceding change may be the more likely cause. Although cross-sectional age-related differences sometimes mirror within-individual longitudinal changes, especially in healthy populations (Salthouse, 2011a), this need not be so (Raz and Lindenberger, 2011), and cross-sectional samples of an underlying longitudinal process may either over- or under-estimate mediation effect sizes (Maxwell & Cole, 2007). While longitudinal studies present their own challenges (Salthouse, 2011b), the cross-sectional nature of the current dataset imposes interpretational limitations, which would benefit from examination in longitudinal cohorts.

Even assuming a causal role of age, questions remain regarding the relationship between neural responsiveness and IQ. Sensitivity analysis indicated robustness to moderate confounders, suggesting a direct effect, although with uncertain direction. We chose to treat IQ as the outcome variable, because this is what we would ultimately hope to improve, and because lesion and neuro-stimulation studies suggest that the integrity and function of frontoparietal networks do causally influence IQ (Glascher et al., 2010; Woolgar et al., 2010; Barbey et al., 2014; Momi et al., 2020; Smith et al., 2022). Although it is hard to imagine experimental manipulations of IQ that are not causally dependent on neural responses, we would not claim that the neural response to cognitive demand is unaffected by IQ. Reduced IQ might either increase MDN responsiveness, if puzzles are experienced as more challenging, or reduce responsiveness, if people confidently select incorrect responses or disengage from the task, making the sign of any reverse relationship difficult to predict. Nonetheless, some combination of reciprocal relationships between IQ and MDN function remains likely, and challenging to disentangle.

Finally, we consider the association between more varied physical activity and attenuation of the dependence of IQ on BOLD responsiveness. While this was consistent across control analyses, the other moderated mediation results should be viewed cautiously as they were affected by covarying head-motion. Again, the nature of the causal links are difficult to determine. First, there are various biological mechanisms by which an active lifestyle might moderate effects of age on the brain, or brain function on cognition (Cotman and Berchtold, 2002; Cotman et al., 2007; Barnes, 2015). Here, the observed moderation altered the second of these links: that is, although engaging in diverse exercise did not reduce the effect of age on the MDN response, it did buffer its cognitive impact, so the reduced MDN response was less strongly associated with reduced IQ. This is consistent with evidence that exercise may support cognition by improving the efficiency of neural function (Neubauer and Fink, 2009; Voss et al., 2011). Indeed, while people with higher IQ may plausibly seek out more activities, better handle a busier routine, or recall more activities, the literature generally supports a causal effect of exercise on cognition (Smith et al., 2010; Liu-Ambrose et al., 2018).

Assuming a causal benefit of physical activity, it is interesting that its variety appears more important than its frequency or duration, consistent with a previous report (Angevaren et al., 2007), and echoing a role of task-novelty in protecting against cognitive decline (Oltmanns et al., 2017). An additional residual effect of exercise duration emerged after covarying head-motion, although this was weaker and less consistent. This finding could provide a principle for designing activity-based interventions with a focus on variety, which may help older adults who are unable or unwilling to perform single, intensive physical activities. Increasing the variety of activities, would represent a lifestyle modification that could be made relatively easily, regardless of specific interests or abilities. Since we examined recent activities, within the preceding year, increasing their variety might be beneficial at any age (cf. Chan et al., 2018).

Interestingly, we did not find that the variety of more intellectual and social activities moderated the same brain∼IQ pathway. This null result could reflect insensitivity of the particular non-physical measures, or a unique role of physical activities in boosting neural efficiency, perhaps via growth-factor-mediated neurogenesis and angiogenesis (Cotman and Berchtold, 2002; Cotman et al., 2007), and consistent with mouse models (Kobilo et al., 2011). Other evidence in humans suggests that the conjunction of physical exercise and mental engagement may especially benefit cognitive function during healthy aging (Fabre et al., 2002). Assuming that monotonous activity is less cognitively stimulating, our measure of variety in exercise may be one way to capture this combination of physical and mental engagement. An effect of non-physical activities *did* emerge after covarying head-motion; however, this moderated the preceding age∼brain pathway so may reflect a different mechanism to varied physical exercise. More non-physical activities were associated with stronger brain responses at younger ages but weaker responses at older ages (crossing at around 50 years). Both observations support previous suggestions that physical and non-physical activities differentially impact late-life cognition, with the latter benefitting more at younger ages (Gow et al., 2017).

In conclusion, we confirm and characterise a neural mechanism that partially explains age-related differences in fluid intelligence, namely reduced responsiveness of the frontoparietal multiple-demand network to cognitive challenge. Specification of such a neuro-cognitive mechanism may facilitate design of targeted interventions to maintain fluid intelligence into healthy old age. As one example, we identify a widely-applicable candidate lifestyle strategy – variety of regular physical activity - that might buffer age-related cognitive decline by decoupling one link in this putative causal pathway.

## Acknowledgements

The Cambridge Centre for Ageing and Neuroscience (Cam-CAN) research was supported by the Biotechnology and Biological Sciences Research Council (grant number BB/H008217/1). Daniel Mitchell and John Duncan were supported by the Medical Research Council intramural program SUAG/085.G116768. We are grateful to the Cam-CAN respondents and their primary care teams in Cambridge for their participation in this study. The original version of the fMRI task was kindly provided by Alexandra Woolgar. The definition of the Core Default Mode network ROI was kindly provided by Moataz Assem.

The Cam-CAN corporate author consists of the Project Principal Personnel: Lorraine K Tyler, Carol Brayne, Edward T Bullmore, Andrew C Calder, Rhodri Cusack, Tim Dalgleish, John Duncan, Richard N Henson, Fiona E Matthews, William D Marslen-Wilson, James B Rowe, Meredith A Shafto; Research Associates: Karen Campbell, Teresa Cheung, Simon Davis, Linda Geerligs, Rogier Kievit, Anna McCarrey, Abdur Mustafa, Darren Price, David Samu, Jason R Taylor, Matthias Treder, Kamen A Tsvetanov, Janna van Belle, Nitin Williams; Research Assistants: Lauren Bates, Tina Emery, Sharon Erzinçlioglu, Andrew Gadie, Sofia Gerbase, Stanimira Georgieva, Claire Hanley, Beth Parkin, David Troy; Affiliated Personnel: Tibor Auer, Marta Correia, Lu Gao, Emma Green, Rafael Henriques; Research Interviewers: Jodie Allen, Gillian Amery, Liana Amunts, Anne Barcroft, Amanda Castle, Cheryl Dias, Jonathan Dowrick, Melissa Fair, Hayley Fisher, Anna Goulding, Adarsh Grewal, Geoff Hale, Andrew Hilton, Frances Johnson, Patricia Johnston, Thea Kavanagh-Williamson, Magdalena Kwasniewska, Alison McMinn, Kim Norman, Jessica Penrose, Fiona Roby, Diane Rowland, John Sargeant, Maggie Squire, Beth Stevens, Aldabra Stoddart, Cheryl Stone, Tracy Thompson, Ozlem Yazlik; and administrative staff: Dan Barnes, Marie Dixon, Jaya Hillman, Joanne Mitchell, Laura Villis, Ethan Knights.

